# aMpLiTuDe MoDuLaTeD noise for tinnitus suppression in tonal and noise-like tinnitus

**DOI:** 10.1101/749937

**Authors:** S. Schoisswohl, J. Arnds, M. Schecklmann, B. Langguth, W. Schlee, P. Neff

## Abstract

**Background:** Acoustic stimulation offers a potential treatment approach for tinnitus but also insights in its basic mechanisms by short-term tinnitus suppression called residual inhibition (RI). The effects of RI were found to be depending on intensity, length or sound types covering the individual tinnitus characteristics. In patients with tonal tinnitus RI was increased with amplitude modulated (AM) pure tones at the individual tinnitus frequency while the effects of modulated noise sounds have not been systematically researched.

**Objectives:** The aim of the present study was to investigate whether in patients with noise-like tinnitus RI can be increased by AM noise-like stimuli according to the individual tinnitus frequency range.

**Methods:** For this purpose the individual tinnitus characteristics (noise-like and tonal tinnitus) were assessed via customizable noise-band matching, in order to generate bandpass filtered stimuli according to the individual tinnitus sound (individualized bandpass filtered sounds; IBP). Subsequent, various stimuli differing in bandpass filtering and AM were tested with respect to their potential to induce RI. Patients were acoustically stimulated with seven different types of stimuli for three minutes each and had to rate the loudness of their tinnitus after each stimuli.

**Results:** Results indicate a general efficacy of noise stimuli for the temporary suppression of tinnitus, but no significant differences between AM and unmodulated IBP. Significantly better effects were observed for the subgroup with noise-like tinnitus (n=14), especially directly after stimulation offset.

**Conclusions:** The study at hand provides further insights in potential mechanisms behind RI for different types of tinnitus. Beyond that, derived principles may qualify for new or extend current tinnitus sound therapies.

## Introduction

Chronic subjective tinnitus is defined as the permanent perception of a sound such as ringing or hissing in the absence of an external or internal source of noise. Approximately 10-15% of the population in industrial countries experience this phantom sound [Langguth et al., 2013; Erlands-son and Dauman, 2013; Heller, 2003; Hall et al., 2011]. Causes for the development of tinnitus are divergent and not completely understood, though most commonly tinnitus occurs towards cochlear damages due to noise trauma [Langguth et al., 2013]. In the majority of cases, the perceived tinnitus pitch is in accordance with the frequency spectrum of hearing loss (HL) [Basile et al., 2013; Roberts et al., 2008]. As a consequence of decreased or absent auditory input and the subsequent deficiency of neural input, maladaptive pathological changes in the auditory pathway are formed, which lead to the perception of a “phantom sound” defined as tinnitus [Eggermont, 2007; Eggermont and Tass, 2015; Eggermont and Roberts, 2012]. Neurophysiological investigations were able to demonstrate hyperactivity in auditory brain areas [Farhadi et al., 2010; Folmer, 2007] as well as aberrant oscillatory brain activity and connectivity patterns [Schlee et al., 2009, 2014; Moazami-Goudarzi et al., 2010; Mohan et al., 2016], in tinnitus patients. Available treatment options for tinnitus have only limited efficacy and to date there is no cure available [Baguley et al., 2013]. Auditory stimulation is one potential treatment approach for tinnitus, but also provides insights to basic mechanisms of tinnitus [Roberts et al., 2008; Fournier et al., 2018].

Almost half a century ago, Feldmann and colleagues investigated the phenomenon of short-term tinnitus suppression after sound stimulation [Feldmann, 1971, 1983]. This temporary suppression is referred to as “residual inhibition” (RI), which manifests in individual suppression patterns (i.e., duration, depth and shape) and can be triggered in 60-80% of subjects with tinnitus [Roberts, 2007; Vernon and Meikle, 2003]. Various recent studies scrutinized RI in more depth. Data from several investigations suggest the effects of RI to be more prominent with sounds close or within the individual tinnitus frequency spectrum [Roberts et al., 2006, 2008; Schaette et al., 2010]. Equally, factors like duration or intensity of the stimuli are essential for its mode of action [Terry et al., 1983; Norena et al., 2002; Vernon and Fenwick, 1984; Neff et al., 2017]. In contrast, the underlying neurophysiological mechanisms of RI are not clearly understood yet [Roberts, 2007; Galazyuk et al., 2019]. Most recent work suggests that tinnitus suppression through sound stimulation is related to reduced spontaneous firing of central auditory neurons [Galazyuk et al., 2017, 2019].

The importance of stimulation intensity and frequency was verified in a recent work from Fournier et al. (2018) [Fournier et al., 2018], who developed a novel approach for RI testing described as Minimum Residual Inhibition Level. Thereby, patients had to adjust the intensity of customized stimuli up to the point where their tinnitus is suppressed during a given interval after the offset of the stimulus. Results show an occurrence of RI in 86.7% of patients by the usage of this method [Fournier et al., 2018].

Despite the manifestation of tinnitus perception as noise-like in many patients, to the best of our knowledge none of the previous mentioned studies included a matching for the band-width of noise-like tinnitus. Those which considered noise-like tinnitus for their methodological approaches, merely used likeliness rating methods for tinnitus matching [Roberts et al., 2006; Fournier et al., 2018].

Recently Henry et al. (2013) [Henry et al., 2013] proposed a novel approach for tinnitus matching procedures taking into consideration the tinnitus type. In addition to the determination of the centre frequency, patients could also adjust the band-width of their tinnitus [Henry et al., 2013]. Here we aim to use both frequency and band-with information to develop individualized stimuli, especially for patients with noise-like tinnitus, for the investigation of residual inhibition.

In previous studies the effects of differently modulated sounds on RI were investigated. These studies revealed that amplitude modulated (AM) tones near or at the individual tinnitus frequency result in larger RI [Reavis et al., 2012; Tyler et al., 2014] with differential results for specific amplitude modulation rates [Neff et al., 2017, 2019].

The hereafter described experiment aims at investigating the effects of different noise stimuli with and without AM on RI. The overarching goal is to establish new acoustic stimulation techniques for basic RI research as well as generating principles for possible future sound stimulation principles with the AM stimulus class. For this purpose, the individual tinnitus characteristics are assessed via noise-band matching as suggested by Henry et al. (2013) [Henry et al., 2013] in order to create personalized stimuli for the RI examination.

Previous studies in the field of RI, already emphasized the impact of noise stimulation on tinnitus perception in tonal tinnitus [Henry et al., 2013; Fournier et al., 2018; Roberts et al., 2006, 2008]. To the best of our knowledge, none of the existing experiments systematically investigated these noise stimulation methods, in particular the application of AM or bandpass filters (BP) to noise stimuli, in noise-like tinnitus.

According to this, the current experiment represents the first attempt to investigate the effects of an administration of individualized BP settings (IBP) and different rates of AM (10 and 40 Hz) to white noise on RI.

These stimulation methods are furthermore merged to a novel combinatory approach to apply IBP and AM to white noise (WN) simultaneously and scrutinize its efficacy in RI.

Additionally each of the used stimuli was examined with regards to induced arousal and valence as rated by the participants.

Besides the assumption of the efficacy of all deployed noise stimuli in short-term tinnitus inhibition (in both noise-like and tonal tinnitus), we expect that IBP differs in its effects on RI from unadjusted WN. Concretely, we presume that the IBP will result in differential residual tinnitus suppression as compared to WN. Yet, given the lack of previous studies we are not able to define a directed hypothesis here. Furthermore, building on the insights of previous work, we hypothesize that stimulations with AM noise (filtered or unchanged) result in larger RI than their unmodulated counterparts.

## Methods

### Participants

The sample for this experiment consisted of N = 29 participants (7 female) between 18 and 75 years with noise-like (n = 14) or tonal tinnitus (n = 15) with a tinnitus duration of more than six months. Participants were recruited from the Interdisciplinary Tinnitus Centre in Regensburg, Germany. For detailed sample characteristics see table 1. Primary inclusion criteria were no somatic, mental or neurological conditions and no current intake of psychotropic medications or substances. Alike, participants were not allowed to participate in other tinnitus-related studies. The methods and the procedures used in this study were examined and approved by the local ethics committee of the University of Regensburg (16-101-0061). All participants were sufficiently informed about the aim, methods and duration of the study, possible side effects, and gave written informed consent prior to the start of the experiment.

**Table 1:**
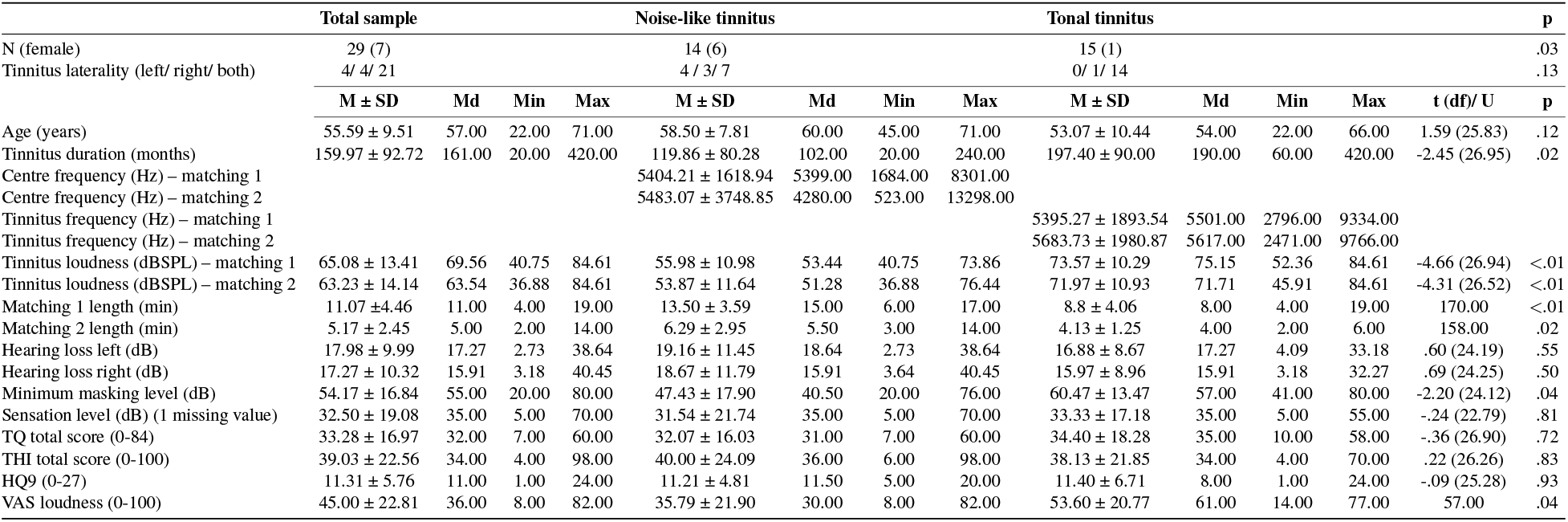
Sample characteristics. M = mean; SD = standard deviation; Md = median; Min = minimum; Max = maximum; TQ = Tinnitus Questionnaire; THI = Tinnitus Handicap Inventory; Mini-HQ9 = Mini Hyperacusis Questionnaire; VAS loudness = Visual Analog Scale tinnitus loudness

### Psychometry

Each participant filled in an online survey composed of German versions of the Tinnitus Handicap Inventory (THI) [Newman et al., 1994; Kleinjung et al., 2007], the Tinnitus Questionnaire (TQ) [Goebel and Hiller, 1994; Hallam et al., 1988], a brief version of the Hyperacusis Questionnaire (mini-HQ9) [Goebel et al., 2013] and the Tinnitus Sample Case History Questionnaire (TSCHQ) for tinnitus-related clinical and demographic information [Langguth et al., 2007].

### Audiometry

For the purpose of individual hearing threshold determination, frequencies ranging from 125 Hz to 8kHz in octave steps including semi-octave steps between 0.5 and 1 (i.e., 0.75 kHz), 1 and 2 (i.e., 1.5 kHz), 2 and 4 (i.e., 3 kHz) and 4 and 8 kHz (i.e., 6 kHz) were quantified with a clinical audiometer (Madsen Midimate 622D; GN Otometrics, Denmark). Sennheiser HDA 2000 headphones (Sennheiser, Germany) were used for audiometric measurements, subsequent tinnitus matching and acoustic stimulation. Minimum Masking Level (MML) was assessed by increasing the loudness of a WN sound (Madsen Midimate 622D; GN Otometrics, Denmark) until their tinnitus was completely masked.

### Tinnitus matching

In order to ascertain participants individual tinnitus pitch, the Method of Adjustment approach (MOA) [Henry et al., 2013] was performed with a custom-made MAX application (MAX 7; Cycling’74, USA) together with a modular hardware controller (Palette Expert Kit; Palette, Canada). The matching procedure’s steps were in accordance with the order within the Tinnitus Tester procedure [Roberts et al., 2008] with an additional test for octave confusion at the end. Prior tinnitus matching, participants were asked to vocalize or describe their tinnitus to distinguish between noise-like and tonal tinnitus types as indicated in the recruiting process. Following on that, they were instructed and trained for the process of tinnitus matching. Parameters examined by the matching procedure were as follows: tinnitus frequency, respectively centre frequency for noise-like tinnitus (Hz), tinnitus loudness (dB) and tinnitus laterality (0 = left ear; 127 = right ear; thus a value of 63 describes a bilateral tinnitus). Control units of the matching controller were labelled accordingly. Step size of frequency dial was marginally below a semitone and ranged from 40 Hz to 16kHz. For tonal tinnitus matching, a 3 kHz pure tone with comfortable loudness was set as a starting point, followed by an adjustment of the frequency by the participants to determine their individual tinnitus frequency. Finally, tinnitus loudness and laterality were adjusted with the matching controller to complete the matching procedure. In case of noise-like tinnitus the starting sound was a filtered broadband noise (bandwidth: 1/3 octave of centre frequency). Patients were able to adjust the centre frequency of the noise and also the bandwidth of the filter settings according to their individual tinnitus noise. Subsequently, loudness and laterality were identified just as with the pure tone matching. Finally, participants rated the agreement of their tinnitus with the matched sound on a 1-10 scale. To assess individuals Sensation Level (SL), the hearing threshold of the frequency next to the individual tinnitus frequency or centre frequency was used (i.e., stepping down to the next lower frequency. e.g., if the individual tinnitus frequency was 7.4 kHz, the hearing threshold at 7 kHz was investigated). The matching procedure was repeated after the acoustic stimulation block of the experiment.

### Acoustic stimulation

Seven different modified noise stimuli were created in MATLAB (Matlab R2015a; Mathworks, USA) and utilised for a three minute acoustic stimulation with an intensity of 60 dB SL. Stimuli set consisted of unmodified WN, WN with AM rates at 10 Hz (WN10) and 40 Hz (WN40), as well as a IBP with the same modulation rates (IBP, IBP10, IBP40). BP width was set according to the matching results in noise-like tinnitus participants. In patients with tonal tinnitus, the previously matched individual tinnitus pitch was used to deploy a IBP to WN with a range of one octave [Pantev et al., 2012]. Furthermore a IBP WN with 10 Hz AM rates at MML intensity (BP10 MML) was used for acoustic stimulation in order to contrast SL and MML. Acoustic stimulation was conducted in a randomized order for each session with a maximum loudness of 80 dBSPL diotically over the headphones. If participants experienced discomfort, they were able to stop the stimulation and experimental procedures at any time. Following a three-minute stimulation for each stimulus, participants evaluated their tinnitus loudness (%) in comparison to prior the particular stimulation on a numeric rating scale (0% up to 140% in 10% steps) at seven different points in time (0, 30, 60, 90, 120, 150 and 180 seconds after stimulation offset). Moreover, participants rated the induced valence and arousal of each single stimuli with pictorial manikin scales [Bradley and Lang, 1994].

### Statistical Analysis

All statistical analysis were performed using the statistic software R (R version 3.4.3; R Foundation for Statistical Computing, Austria) and the packages “psych”, “emmeans”, “sjstats” and “lme4”. Tinnitus loudness and stimulus evaluation (valence and arousal) data were analyzed by means of linear mixed effect models according to the following formula: *Y*_*i*_ ∼ *X*_*i*_*β* + *Z*_*i*_*u*_*i*_ + *ϵ*_*i*_, whereby *Y*_*i*_ represents the dependent variable, *X*_*i*_ is the particular predictor or so called fixed effect of the model with *β* as its weight estimates. The notion *Z*_*i*_ describes the random effect with the corresponding random vector *u*_*i*_, plus *ϵ*_*i*_ serves as the random vector of the model fit error. In order to identify the respective model with the best fit for the data, a step-wise selection approach was carried out by gradually adding a new fixed effect to the model. In a next step the model was compared to a corresponding “null” model without the fixed effect with a Likelihood Ratio Test (LRT) [Harrison et al., 2018]. Model-fitting procedure was performed for each dependent variable, denoted as response (tinnitus loudness, valence, arousal), individually and tested the following predictors as well as their interactions: condition (stimuli used; see acoustic stimulation section), group (noise-like tinnitus, tonal tinnitus), time (0sec, 30sec, 60sec, 90sec, 120sec, 150sec, 180sec after stimulation end), gender (male, female), age, tinnitus duration, tinnitus loudness (according to first tinnitus matching), MML and tinnitus distress (TQ sum score). The proportion of explained variance was identified by marginal (variance of the fixed effects) and conditional (variance of fixed and random effects) *R*^2^ [Nakagawa et al., 2017]. In any of the fitted models, participant (id) was treated as a random effect. Fixed effects of the final model were tested via expected mean square approach. Post-hoc Tukey-tests were calculated to contrast responses for condition and group. In order to test for a potential bias due to the sequence of the stimuli used for acoustic stimulation (position effect), a median split was conducted on the positions variable and differences in means were then tested with student t-tests.

Analysis of descriptive group differences (noise-like vs. tonal tinnitus) for parametric variables were conducted by the means of two-sample t-tests. Assumptions of normal distribution (Shapiro-Wilk-Test) and homoscedasticity (F-test) were tested and if violated, non-parametric testing via independent sample Mann-Whitney U-tests was used.

Categorical data was analyzed by Fisher’s exact tests, due to cell frequencies below 5 in all variables.

Reliability for the matching procedure (between first and second matching round) was assessed via Pearson correlations, or rather Spearman correlations in case of a violation of normal distribution, for tinnitus loudness and tinnitus or centre frequency. Statistical significance was defined as *p* ≤ .05 for all analysis.

## Results

### Descriptives

Demographic and clinical characteristics for the whole study sample and for tinnitus subgroups (noise-like and tonal tinnitus) can be found in table 1. A Fisher’s exact test was able to identify a significant association between gender and the type of tinnitus. In the group with tonal tinnitus the proportion of female patients was significantly lower (p = .03). Statistical testing revealed significant differences in terms of tinnitus duration and the subjective rating of tinnitus loudness (VAS loudness), with noise-like tinnitus patients showing a shorter duration of tinnitus (t _(26.95)_ = −2.45, p = .02) and evaluating their tinnitus loudness lower (U = 57.00, p = .04). Further, no differences were found in TQ (t _(26.90)_ = −.36, p = .72), THI (t _(26.26)_ = .22, p = .83) or HQ9 (t _(25.28)_ = −.09, p = .93) scores among the two subgroups.

### Audiometry and Tinnitometry

Table 1 shows audiometric and tinnitus matching results with a significant lower tinnitus loudness (corresponding with subjective loudness rating; see descriptives section above) for both matching procedures (matching 1: t _(26.94)_ = −4.66, p *<* .01; matching 2: t _(26.52)_ = −4.31, p *<* .01) and MML t _(24.12)_ = −2.20, p = .04) in the group of noise-like tinnitus. On the basis of a consolidation of these audiometric and tinnitometric findings, figure 1 indicates an overlap of tinnitus frequency with the frequency of HL. As might be expected, the length of the first and second matching process was significant shorter in tonal tinnitus patients (cf. table 1). Mean HL difference for both ears were not significant different between groups (left: t _(24.19)_ = .60, p = .55; right: t _(24.25)_ = .69, p = .50). In both groups the HL was more pronounced on the left side.

**Figure 1:**
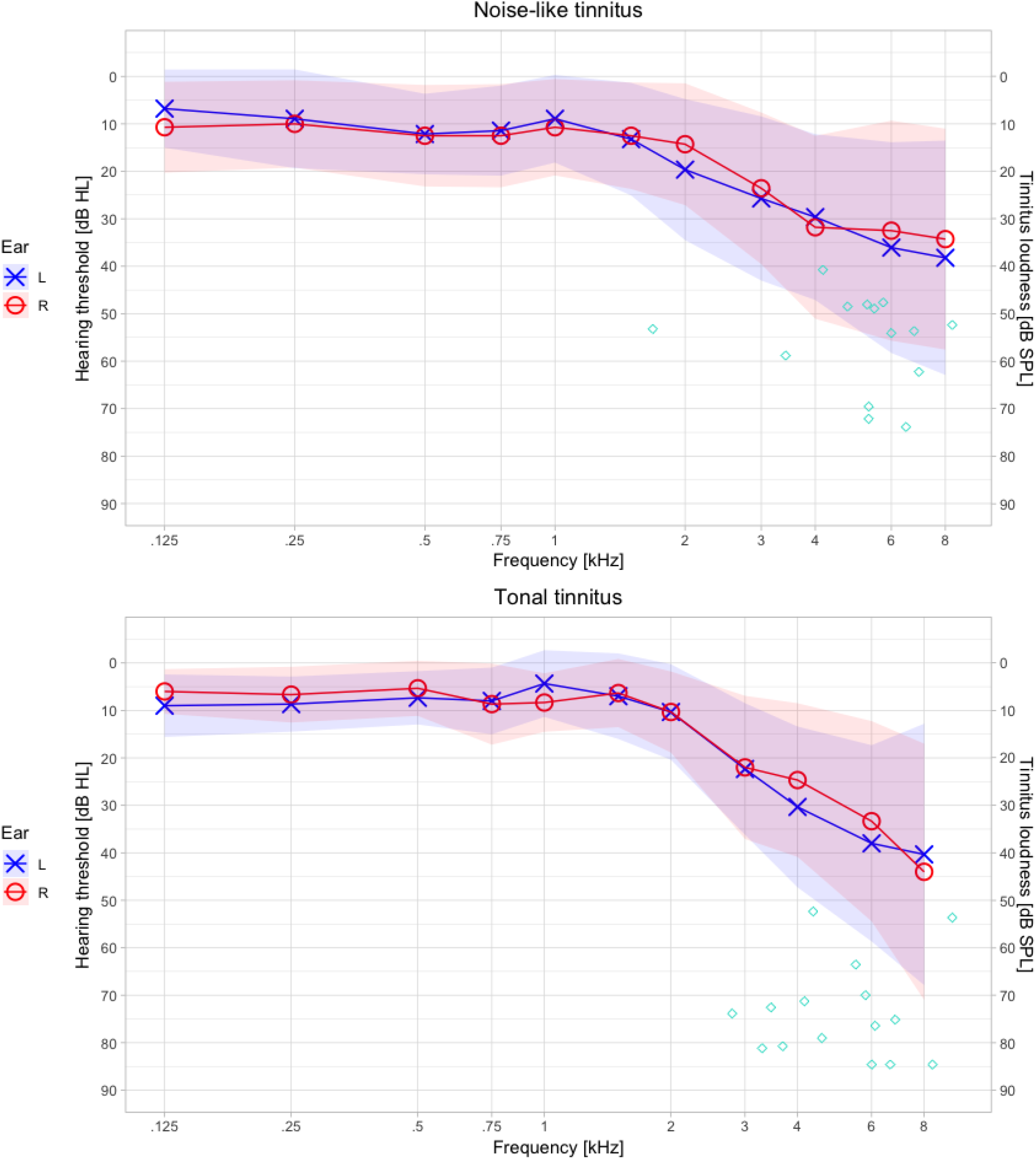
Audiometry and Tinnitometry. Audiometric measurement results for both ears together with individual tinnitus frequency (i.e., centre frequency of the IBP) and loudness as identified by tinnitus matching splitted for noise-like and tonal tinnitus. It should be noted, that tinnitus/ centre frequency overlaps with the frequencies of hearing loss.

There were positive significant correlations between the first and the second matching for tinnitus loudness (noise-like: r = .77, p *<* .01; tonal: r = .73, p = *<* .01) in both groups. With respect to tinnitus/ centre frequency a positive significant correlation was only observed in the tonal tinnitus group (noise-like: r = .14, p = .64; tonal: r = .65, p = *<* .01).

### Acoustic stimulation

Prima facie, the stimulus IBP40 appeared to produce the strongest tinnitus suppression regardless of group and time (M = 86.16, SD = 25.60), whereas at timepoint T0 (immediately after stimulation offset), WN40 induced the lowest tinnitus loudness (M = 73.10, SD = 41.76). Descriptive statistics for the 7 utilized stimuli averaged over time and for timepoint T0 are listed in table S1 for the whole sample and divided for subgroups. Figure 2 shows the time curve for all stimuli with respect to tinnitus loudness ratings, in the same manner figure S1 provides information about single subject responses for each stimuli. No confounding effect caused by the order of the stimuli in the stimulation sequence was detected by our analysis (t_(1215.60)_ =.09, p = .93) and therefore position was not entered in the final model fitting procedure. In accordance with the previous described model fitting approach (cf. section statistical analysis in methods part), we were able to identify the following model with the best fit to our data: *response ∼ condition* + *time ∗ group* + (1 *| id*). Detailed results of the model fitting are outlined in table S2. By testing the fixed effects of the model via expected mean square approach, significant effects for condition, time, group and for the interaction time*group on tinnitus loudness were observed (cf. table 2). Subsequent post-hoc contrasts for condition failed to find statistically significant differences in tinnitus loudness ratings with respect to the applied stimuli (see table 3). Interestingly, a significant difference in tinnitus loudness ratings between the two subgroups was revealed independently of condition and time as exemplified in table 4 and figure 3 (noise-like: M = 82.14, SD = 26.68; tonal: M = 94.79, SD = 16.44; t_(31.15)_ = 2.17, p = .04). On the basis of a significant interaction among group and time, we contrasted the mean tinnitus loudness for each group for all 7 timepoints after stimulation. Our results point out a significant difference between the groups only at T0 (noise-like: M = 63.98, SD = 36.49; tonal: M = 90.19, SD = 28.01; t_(38.40)_ = 4.27, p *<* .01) (cf. table 5).

**Table 2:** Fixed effect testing. numDF = degrees of freedom numerator; denDF = degrees of freedom denominator

|  | numDF | denDF | F | p |
| --- | --- | --- | --- | --- |
| Condition | 6.00 | 1392.00 | 3.35 | <.01 |
| Time | 6.00 | 1392.00 | 39.84 | <.01 |
| Group | 1.00 | 29.00 | 5.04 | .03 |
| Time*Group | 6.00 | 1392.00 | 15.17 | <.01 |

**Table 3:** Post-hoc tukey contrasts for condition. Degrees of freedom = 1410.23; standard error = 1.44

| **Contrast** | **Estimate** | **t** | **p** |
| --- | --- | --- | --- |
| IBP - IBP10 | -1.53 | -1.06 | 0.94 |
| IBP - IBP10.MML | -4.38 | -3.05 | 0.04 |
| IBP - IBP40 | 1.08 | 0.75 | 0.99 |
| IBP - WN | -2.76 | -1.92 | 0.47 |
| IBP - WN10 | -2.17 | -1.51 | 0.74 |
| IBP - WN40 | -0.34 | -0.24 | >.99 |
| IBP10 - IBP10.MML | -2.86 | -1.98 | 0.42 |
| IBP10 - IBP40 | 2.61 | 1.81 | 0.54 |
| IBP10 - WN | -1.23 | -0.86 | 0.98 |
| IBP10 - WN10 | -0.64 | -0.44 | >.99 |
| IBP10 - WN40 | 1.18 | 0.82 | 0.98 |
| IBP10.MML - IBP40 | 5.47 | 3.80 | <.01 |
| IBP10.MML - WN | 1.63 | 1.13 | 0.92 |
| IBP10.MML - WN10 | 2.22 | 1.54 | 0.72 |
| IBP10.MML - WN40 | 4.04 | 2.81 | 0.08 |
| IBP40 - WN | -3.84 | -2.67 | 0.11 |
| IBP40 - WN10 | -3.25 | -2.26 | 0.27 |
| IBP40 - WN40 | -1.43 | -0.99 | 0.96 |
| WN - WN10 | 0.59 | 0.41 | >.99 |
| WN - WN40 | 2.41 | 1.68 | 0.63 |
| WN10 - WN40 | 1.82 | 1.27 | 0.87 |

**Table 4:**
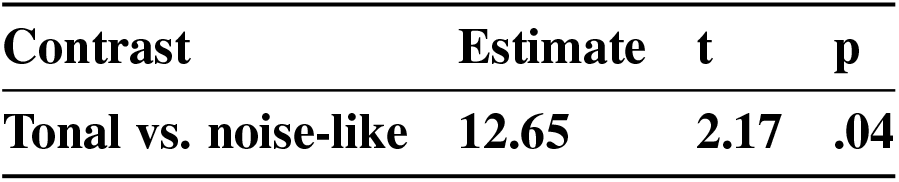
Post-hoc tukey contrasts for group. Degrees of freedom = 31.15; standard error = 5.84

| **Contrast** | **Estimate** | **t** | **p** |
| --- | --- | --- | --- |
| **Tonal vs. noise-like** | **12.65** | **2.17** | **.04** |

**Table 5:**
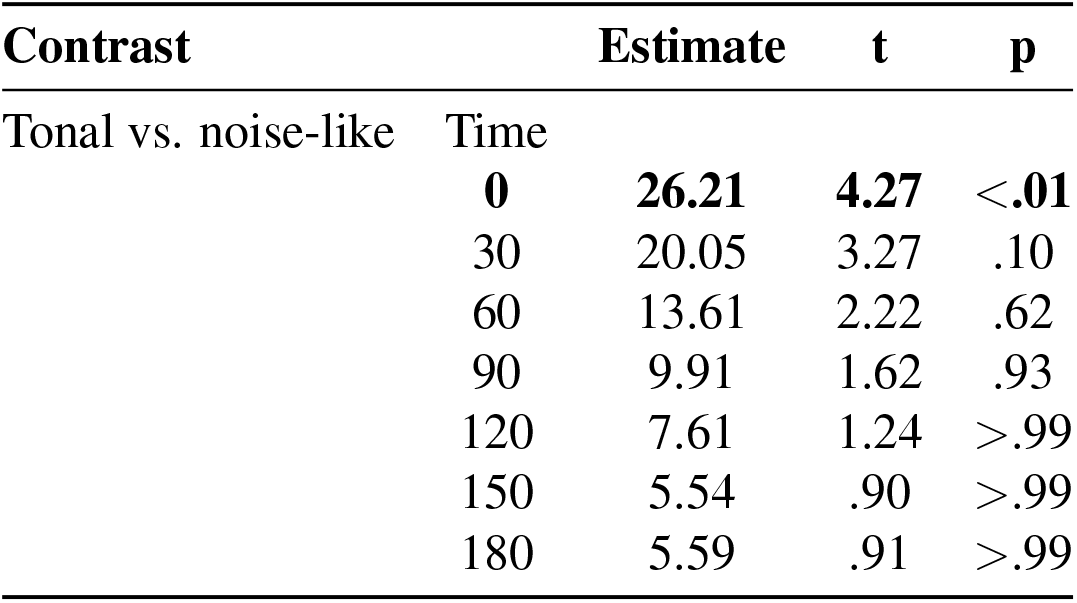
Post-hoc tukey contrasts for group*time. Degrees of freedom = 38.40; standard error = 6.13

**Figure 2:**
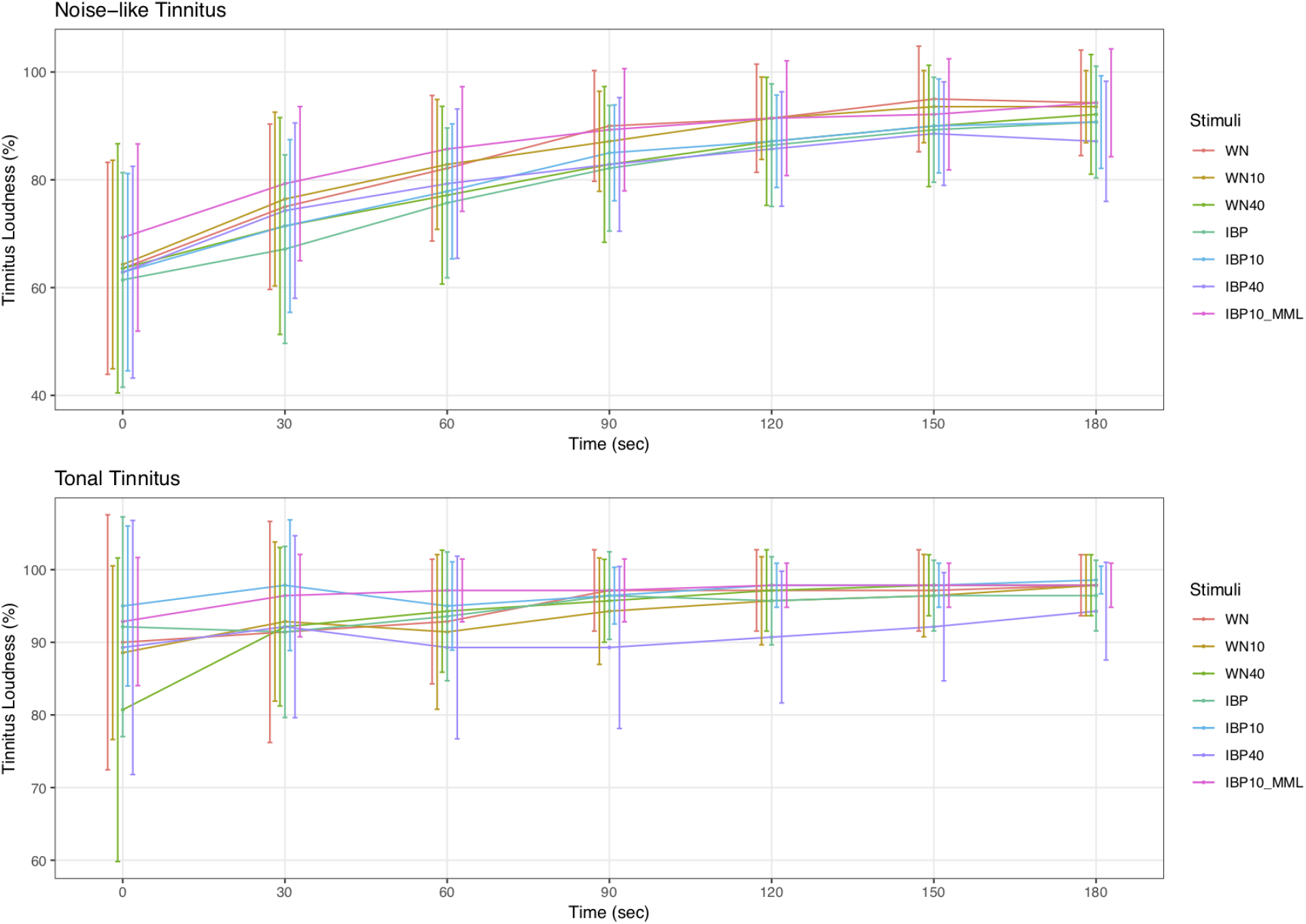
Tinnitus loudness time curve splitted by group. For each stimuli the tinnitus loudness rating over all time points is plotted separated for noise-like and tonal tinnitus (confidence intervals at 95% shown as brackets). Overall, each stimulus was able to suppress tinnitus loudness (cf. table S1). In terms of suppression averaged over time but also at T0, stimulus IBP appeared to produce the strongest effect on loudness in the noise-like tinnitus group. Whereas in the tonal group, stimulus IBP40 induced the lowest tinnitus loudness on average. However, directly after stimulation WN40 showed the strongest suppression.

**Figure 3:**
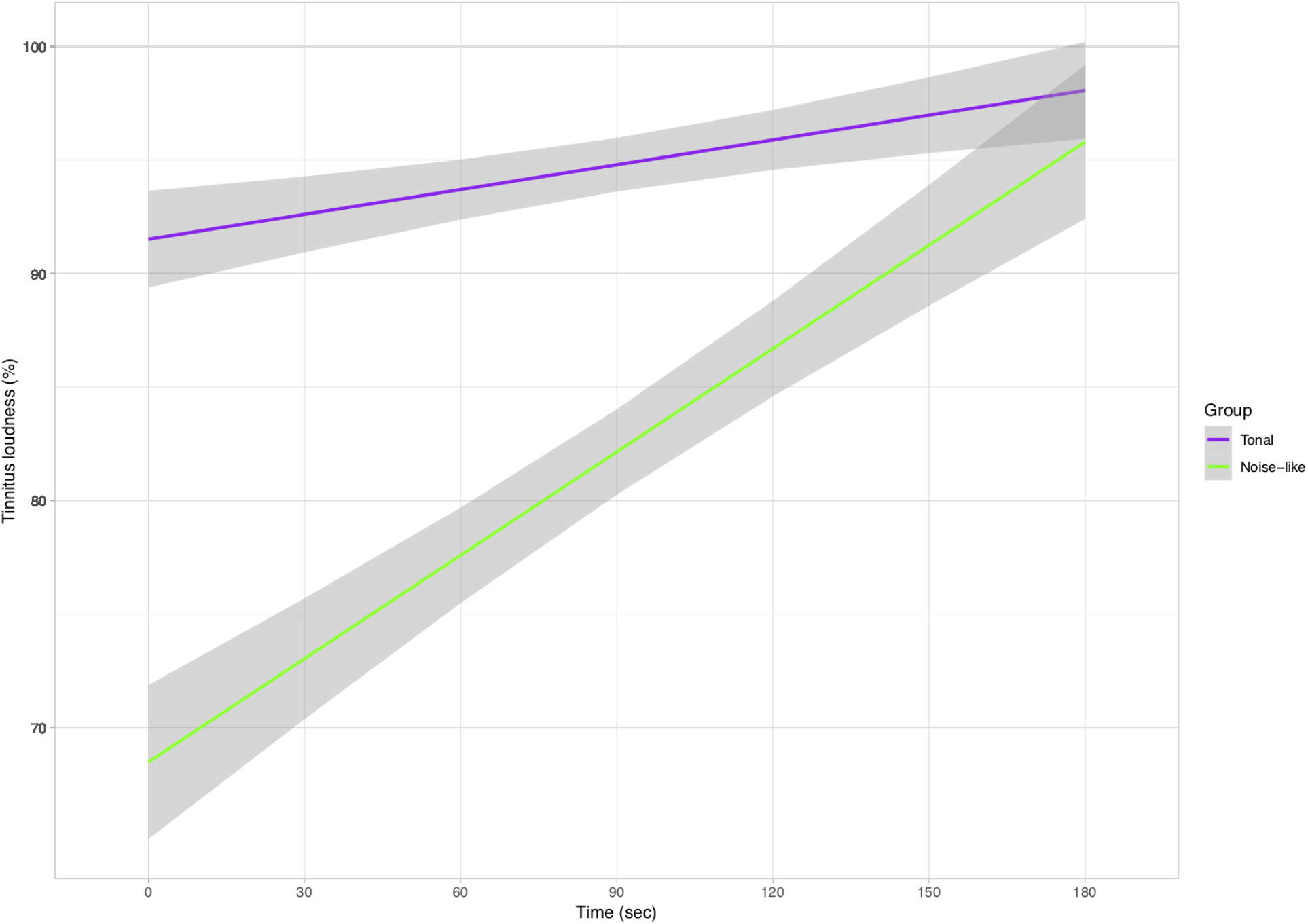
Mean suppression differences between groups. Time curve of the averaged tinnitus suppression values splitted for tonal and noise-like tinnitus. Standard deviation for the mean suppression data of each group is plotted as a grey ribbon. Differences between the two subgroups were found to be significant.

## Stimulus evaluation

### Arousal

As pointed out in table S3 and figure 4, emotional stimuli evaluation for the whole group identified the highest arousal ratings for stimulus IBP40, while IBP10 MML expectably manifested in the lowest arousal values. Model fitting proceedings identified the subsequent model with the best fit for our arousal data: *response ∼ condition* + (1 *| id*) (cf. table S4). Fixed effect testing detected a significant effect for condition (cf. table 6). Ensuing post-hoc contrasts revealed significant differences in arousal ratings for IBP vs. IBP40 (t_(180.21)_ = −3.08, p = .04), IBP10 vs. IBP10 MML (t_(180.21)_ = 2.98, p = .05), IBP10 MML vs. IBP40 (t_(180.21)_ = −4.33, p *<* .01), IBP10 MML vs. WN10 (t_(180.21)_ = −3.66, p *<* .01) and IBP10 MML vs. WN40 (t_(180.21)_ = −4.04, p *<* .01). Post-hoc analysis results are reported in table 7, relevant significant results are highlighted in bold.

**Table 6:** Fixed effect testing - arousal & valence. numDF = degrees of freedom numerator; denDF = degrees of freedom denominator

|  | numDF | denDF | F | p |
| --- | --- | --- | --- | --- |
| **Arousal** |  |  |  |  |
| Condition | 6.00 | 174.00 | 5.17 | <.01 |
| **Valence** |  |  |  |  |
| Condition | 6.00 | 174.00 | 3.25 | <.01 |

**Table 7:** Post-hoc tukey contrasts for condition. Arousal: Degrees of freedom = 180.21; standard error = .36; Valence: Degrees of freedom = 180.21; standard error = .45

| Contrast | Arousal |  |  | Valence |  |  |
| --- | --- | --- | --- | --- | --- | --- |
|  | Estimate | t | p | Estimate | t | p |
| IBP - IBP10 | -0.62 | -1.73 | 0.60 | 0.17 | 0.39 | >.99 |
| IBP - IBP10_MML | 0.45 | 1.25 | 0.87 | -0.48 | -1.08 | 0.93 |
| **IBP - IBP40** | **-1.10** | **-3.08** | **0.04** | 0.59 | 1.31 | 0.85 |
| IBP - WN | -0.38 | -1.06 | 0.94 | 0.14 | 0.31 | >.99 |
| IBP - WN10 | -0.86 | -2.41 | 0.20 | 0.79 | 1.77 | 0.57 |
| IBP - WN40 | -1.00 | -2.79 | 0.08 | 1.21 | 2.70 | 0.10 |
| IBP10 - IBP10_MML | 1.07 | 2.98 | 0.05 | -0.66 | -1.47 | 0.76 |
| IBP10 - IBP40 | -0.48 | -1.35 | 0.83 | 0.41 | 0.93 | 0.97 |
| IBP10 - WN | 0.24 | 0.67 | 0.99 | -0.03 | -0.08 | >.99 |
| IBP10 - WN10 | -0.24 | -0.67 | 0.99 | 0.62 | 1.39 | 0.81 |
| IBP10 - WN40 | -0.38 | -1.06 | 0.94 | 1.03 | 2.32 | 0.24 |
| IBP10_MML - IBP40 | -1.55 | -4.33 | <.01 | 1.07 | 2.39 | 0.21 |
| IBP10_MML - WN | -0.83 | -2.31 | 0.25 | 0.62 | 1.39 | 0.81 |
| IBP10_MML - WN10 | -1.31 | -3.66 | 0.01 | 1.28 | 2.86 | 0.07 |
| IBP10_MML - WN40 | -1.45 | -4.04 | <.01 | 1.69 | 3.78 | <.01 |
| IBP40 - WN | 0.72 | 2.02 | 0.41 | -0.45 | -1.00 | 0.95 |
| IBP40 - WN10 | 0.24 | 0.67 | 0.99 | 0.21 | 0.46 | >.99 |
| IBP40 - WN40 | 0.10 | 0.29 | >.99 | 0.62 | 1.39 | 0.81 |
| WN - WN10 | -0.48 | -1.35 | 0.83 | 0.66 | 1.47 | 0.76 |
| WN - WN40 | -0.62 | -1.73 | 0.60 | 1.07 | 2.39 | 0.21 |
| WN10 - WN40 | -0.14 | -0.38 | >.99 | 0.41 | 0.93 | 0.97 |

**Figure 4:**
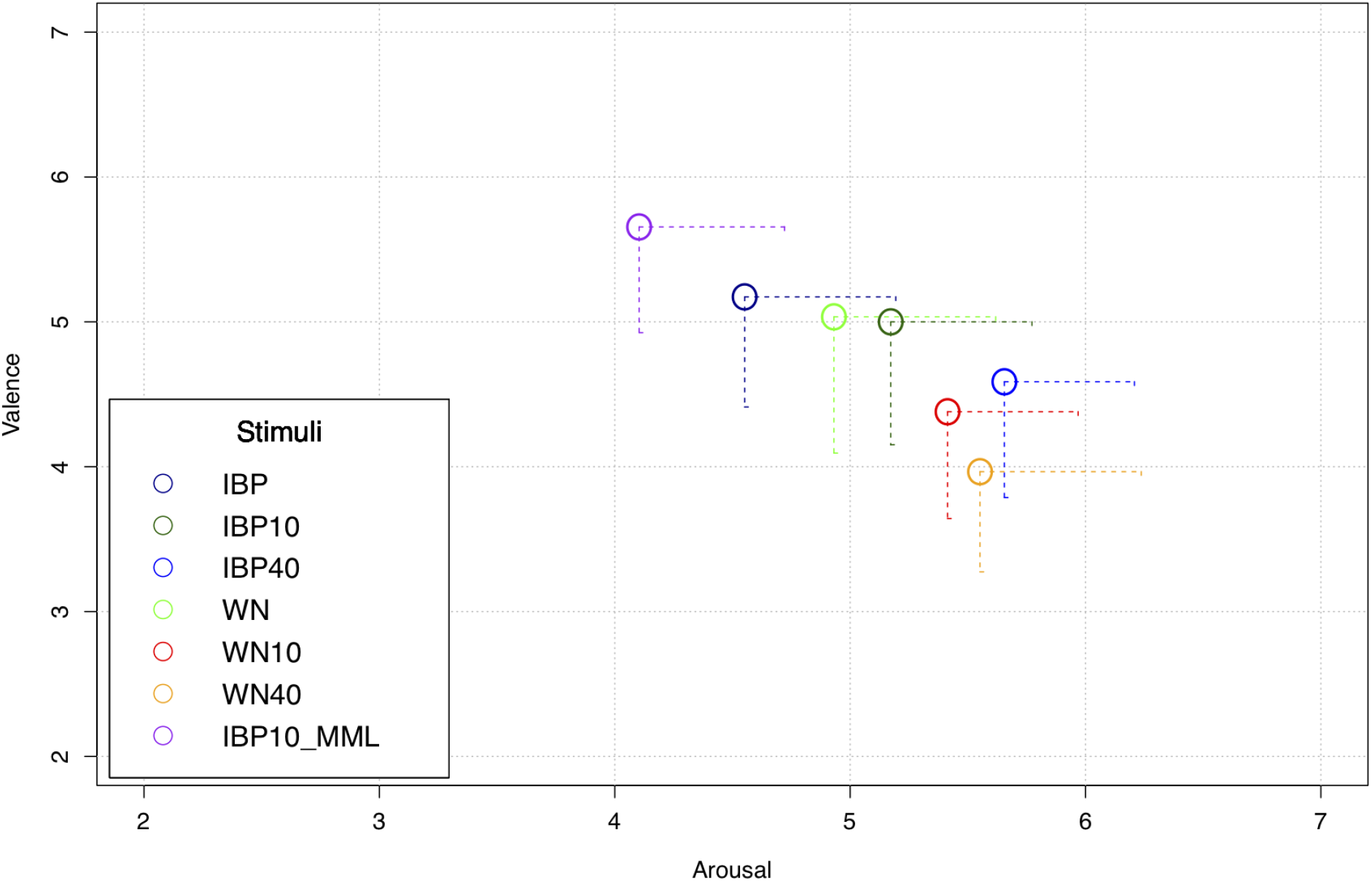
Valence and arousal rating per stimuli. Parentheses show 95 % confidence interval for arousal and valence ratings for all stimuli. Lowest tolerability was found in WN40 as indicated by high arousal and low valence stimulus valuation. Whereas stimulus IBP10 MML shows the highest tolerability.

### Valence

In line with the descriptive arousal results, IBP10 MML had the highest ratings for valence, whereas stimuli WN40 was evaluated with the least valence (cf. table S3 and figure 4). Same model structure was fitted as for the arousal data (cf. table S4) and likewise a significant effect of condition was found (cf. table 6). Post-hoc results are listed in table 7 and demonstrate a significant difference for IBP10 MML vs. WN40 (t_(180.21)_ = 3.78, p *<* .01).

## Discussion

The aim of the present study was to investigate the effects of different IBP and AM noise stimuli on RI in patients with tonal and noise-like tinnitus. To the best of our knowledge, no former study has systematically investigated the deployed acoustic stimulation procedures, especially neither AM or IBP sounds in patients with noise-like tinnitus. A parametric noise-band matching approach was applied in order to personalize BP settings in accordance with the tinnitus characteristics in the group with noise-like tinnitus, whereas the group with tonal tinnitus matched their tinnitus via the centre frequency of a fixed filter bandwidth. Taken together, all these aspects constitute novel lines of investigation within tinnitus research. Omnibus results of our experiment emphasize the ability of all used noise stimuli in inducing RI (cf. table 2). The time courses and different suppression patterns for each stimuli appear in a similar manner as in previous studies, in that they generally converge over time after an initial maximum of suppression [Feldmann, 1983; Roberts et al., 2008; Neff et al., 2017, 2019; Vernon and Meikle, 2003; Roberts, 2007].

Contrary to our hypotheses, the central finding of our analysis indicates no statistically significant differences between the various stimuli and their impact on tinnitus perception respectively RI. In more detail, neither the customization of the noise bands nor the AM resulted in significant differences between the conditions (i.e., stimuli). This outcome is in conflict with findings of earlier studies, which have suggested advantages of AM pure tones for RI [Neff et al., 2017, 2019; Reavis et al., 2012; Tyler et al., 2014]. Yet, looking at these studies, merely pure tones were compared to AM pendants with the exception of Tyler et al. (2014) [Tyler et al., 2014], who contrasted AM pure tones with unmodulated broadband noise. No former study aimed at investigating AM and IBP noises for RI or sound therapy, especially in noise-like tinnitus, which renders the discussion of the current results difficult. These observations, while not explaining the non-existing effects in this study, certainly help to better understand the parameters of RI stimuli (here: carrier sounds and modulation rates) in the research branch of acoustic stimulation in tinnitus. Alternatively, a potential explanation for the lack of advantages of AM stimuli could be attributed to the circumstances, that noise is inherently composed of a wide spectrum of frequencies and signal-inherent amplitude modulation rates. These may cover up or neutralize the potential effects of certain AM rates for RI.

To the best of our knowledge, no former study specifically tested RI or sound therapies in patients with noise-like tinnitus. Of special interest, our analysis revealed statistical differences in RI for the subgroups noise-like and tonal tinnitus, with noise-like patients demonstrating larger RI than the tonal group. These significant differences were only observed immediately after the stimulation, suggesting a time-limited advantage of noise stimuli for RI in noise-like tinnitus. The reason for this group-difference is not clear, a possible rationale may be due to physiological differences between these two groups with a supposed additional contribution of the extralemniscal system in noise-like tinnitus [Møller, 2006].

A further potential confounding factor for this group effect might be the fact that tinnitus loudness as elicited by MML, tinnitus matching and also in subjective ratings via VAS scales was found to be significant higher in the tonal subgroup. On the other hand, with no meaningful difference in HL between the groups and in consequence similar SLs, the putative confounding influence of these measures may play a negligible role. An in-depth analysis of the noise-like tinnitus group exclusively, demonstrated no statistical differences in tinnitus loudness ratings with respect to the used stimuli in a similar fashion as the analysis of the whole study sample.

However, since bandwidth of BP filter settings in tonal tinnitus patients was set to a range of one octave around the indiviual tinnitus frequency, whereas noise-like patients were able to individually adjust the BP filter settings, the differences in the subgroups may also derive from discrepancies in stimuli creation.

It is naturally supposed, that a stimulation with noise is more pleasant or tolerable than a stimulation with pure tones. Unlike this assumption, our findings reveal a similar tolerability pattern for AM noise stimuli as Neff et al. (2019) [Neff et al., 2019] on the basis of AM pure tones (cf. figure 4). The analysis conducted also show, that AM might lead to more arousal as indicated on a descriptive level as well as the significant difference between IBP and IBP40 (cf. table 7). As must be expected, the lower intensity stimulus (IBP10 MML) had the lowest arousal and highest valence ratings.

Our results indicate that the used matching method is feasible for determining tinnitus characteristics. In detail there was good consistency for both tinnitus loudness and frequency for both matching trials in noise-like and tonal tinnitus groups. These findings are in line with Henry et al. (2013) [Henry et al., 2013], who already reported test-retest reliability for noise-band tinnitus matching.

### Limitations

The generalizability of these results is subject to certain limitations. As already discussed above, the significantly lower tinnitus loudness in the group of noise-like tinnitus could weaken our findings of subgroup differences in short-term tinnitus suppression. However, as no difference in HL and equality in SL were observed, this may not play a significant role.

Likewise, the sample size of this experiment is rather small and gender ratio in the subgroups is unbalanced. One main issue is the impossibility to control for potential participant-related failures in noise-band matching. But for all of that, unavailable validation of the quantification of patients tinnitus characteristics represents a common problem in tinnitus matching approaches, as it is a subjective phenomenon. Future studies should strive for new possibilities in verifying tinnitus matching results, as well as optimization of given methodological approaches.

Due to a lack of tonal stimuli in the present experiment and the missing comparison of tonal and noise stimuli, it is not possible to make a statement about a general superiority of noise stimuli in short-term tinnitus suppression in noise-like tinnitus patients.

## Conclusion

The current study demonstrates a general efficacy of noise stimuli with different AM rates and filtering strategies for RI. Contrary to our expectations, no differences between the types of stimuli were observed, namely between unfiltered WN and IBP as well as unmodulated WN and different AM rates, respectively. Rather, differences in RI among the subgroups of noise-like and tonal tinnitus, with better performance directly after the stimulation in the noise-like tinnitus group, were observed. Although, no stable rationale for the group differences can be provided, the findings may provide insights in the mechanism of RI for different tinnitus types. Future studies with larger sample sizes, improved matching/ audiometry procedures and more acoustic stimulation repetitions per stimuli are needed to investigate these potential differences in more detail in order to enhance our understanding of the effects of acoustic stimulation on tinnitus perception.

Taken together these results illustrate the potential of noise-stimuli in short-term tinnitus suppression, especially in patients with noise-like tinnitus.

## Acknowledgement

We want to thank Susanne Staudinger for her extremely valuable support in the collection of the data. This project was conducted as part of the European School for Interdisciplinary Tinnitus Research (ESIT) [Schlee et al., 2018].

## Statement of Ethics

This study was approved by the ethics committee of the University of Regensburg, Germany (16-101-0061).

## Author Contributions

The authors P.N., W.S and S.S. designed the study. J.A. collected the data. S.S. and P.N. analyzed the data and wrote the main manuscript. All authors contributed to and reviewed the manuscript.

## Supplemental material

**Table S1:** Tinnitus loudness per condition. M = mean; SD = standard deviation; Md = median; Min = minimum; Max = maximum; T0 = immediately after stimulation offset

|  | Total |  |  |  | T0 |  |  |  |
| --- | --- | --- | --- | --- | --- | --- | --- | --- |
|  | M ± SD | Md | Min | Max | M ± SD | Md | Min | Max |
| IBP | 87.24 ± 23.93 | 100.00 | .00 | 120.00 | 77.59 ± 36.22 | 100.00 | .00 | 120.00 |
| IBP10 | 88.77 ± 20.82 | 100.00 | 10.00 | 140.00 | 77.95 ± 32.56 | 80.00 | 10.00 | 140.00 |
| IBP10_MML | 91.63 ± 19.01 | 100.00 | 10.00 | 110.00 | 81.72 ± 28.17 | 100.00 | 10.00 | 110.00 |
| IBP40 | 86.16 ± 24.60 | 100.00 | .00 | 130.00 | 77.59 ± 37.57 | 100.00 | .00 | 130.00 |
| WN | 90.00 ± 23.29 | 100.00 | .00 | 140.00 | 77.59 ± 37.00 | 100.00 | .00 | 140.00 |
| WN10 | 89.41 ± 20.93 | 100.00 | .00 | 120.00 | 77.24 ± 32.28 | 80.00 | .00 | 120.00 |
| WN40 | 87.95 ± 26.51 | 100.00 | .00 | 130.00 | 73.10 ± 41.76 | 100.00 | .00 | 130.00 |
| **Noise-like** |  |  |  |  |  |  |  |  |
| IBP | 78.98 ± 27.86 | 90.00 | .00 | 120.00 | 61.43 ± 38.00 | 65.00 | .00 | 120.00 |
| IBP10 | 80.71 ± 24.67 | 90.00 | 10.00 | 120.00 | 62.86 ± 34.96 | 50.00 | 10.00 | 120.00 |
| IBP10_MML | 85.92 ± 24.49 | 100.00 | 10.00 | 110.00 | 69.29 ± 33.16 | 80.00 | 10.00 | 110.00 |
| IBP40 | 80.10 ± 26.81 | 90.00 | .00 | 120.00 | 62.86 ± 37.50 | 55.00 | .00 | 120.00 |
| WN | 84.49 ± 26.56 | 100.00 | .00 | 110.00 | 63.57 ± 37.54 | 70.00 | .00 | 110.00 |
| WN10 | 84.18 ± 24.41 | 100.00 | .00 | 120.00 | 64.29 ± 36.94 | 65.00 | .00 | 120.00 |
| WN40 | 80.61 ± 31.29 | 100.00 | .00 | 120.00 | 63.57 ± 44.13 | 70.00 | .00 | 120.00 |
| **Tonal** |  |  |  |  |  |  |  |  |
| IBP | 94.95 ± 16.24 | 100.00 | 10.00 | 120.00 | 92.67 ± 27.89 | 100.00 | 10.00 | 120.00 |
| IBP10 | 96.29 ± 12.50 | 100.00 | 50.00 | 140.00 | 92.00 ± 23.36 | 100.00 | 50.00 | 140.00 |
| IBP10_MML | 96.95 ± 9.11 | 100.00 | 50.00 | 110.00 | 93.33 ± 16.33 | 100.00 | 50.00 | 110.00 |
| IBP40 | 91.81 ± 20.93 | 100.00 | 10.00 | 130.00 | 91.33 ± 33.14 | 100.00 | 10.00 | 130.00 |
| WN | 95.14 ± 18.46 | 100.00 | .00 | 140.00 | 90.67 ± 32.40 | 100.00 | .00 | 140.00 |
| WN10 | 94.29 ± 15.68 | 100.00 | 40.00 | 120.00 | 89.33 ± 22.19 | 100.00 | 40.00 | 120.00 |
| WN40 | 94.10 ± 19.05 | 100.00 | .00 | 130.00 | 82.00 ± 38.77 | 100.00 | .00 | 130.00 |

**Table S2:** Model fitting. df = degrees of freedom; AIC = Akaike Information Criterion; BIC = Bayesian Information Criterion; logLik = log-likelihood; LRT = Likelihood Ratio Test

|  | R<sup>2</sup> (marginal) | R<sup>2</sup> (conditional) | df | AIC | BIC | logLik | LRT | p |
| --- | --- | --- | --- | --- | --- | --- | --- | --- |
| Intercept only: response ~ 1 + (1\|id) | .00 | .51 | 3 | 12046.00 | 12061.00 | -6019.00 |  |  |
| Fitted model : response ~ condition + time * group + (1\|id) | .17 | .60 | 22 | 11774.00 | 11890.00 | -5865.20 | 309.22 | <.01 |

**Table S3:** Stimulus evaluation. M = mean; SD = standard deviation; Md = median; Min = minimum; Max = maximum

|  | Arousal |  |  |  | Valence |  |  |  |
| --- | --- | --- | --- | --- | --- | --- | --- | --- |
|  | M ± SD | Md | Min | Max | M ± SD | Md | Min | Max |
| IBP | 4.55 ± 1.76 | 5.00 | 1.00 | 7.00 | 5.17 ± 2.09 | 5.00 | 2.00 | 9.00 |
| IBP10 | 5.17 ± 1.65 | 5.00 | 2.00 | 8.00 | 5.00 ± 2.33 | 5.00 | 1.00 | 9.00 |
| IBP10_MML | 4.10 ± 1.70 | 4.00 | .00 | 8.00 | 5.66 ± 2.00 | 5.00 | 2.00 | 9.00 |
| IBP40 | 5.66 ± 1.52 | 6.00 | 2.00 | 8.00 | 4.59 ± 2.20 | 4.00 | 1.00 | 9.00 |
| WN | 4.93 ± 1.89 | 5.00 | 1.00 | 8.00 | 5.03 ± 2.58 | 5.00 | 1.00 | 9.00 |
| WN10 | 5.41 ± 1.52 | 5.00 | 3.00 | 9.00 | 4.38 ± 2.03 | 4.00 | 1.00 | 9.00 |
| WN40 | 5.55 ± 1.88 | 5.00 | 1.00 | 9.00 | 3.97 ± 1.90 | 3.00 | 1.00 | 9.00 |
| **Noise-like** |  |  |  |  |  |  |  |  |
| IBP | 3.86 ± 1.88 | 4.00 | 1.00 | 7.00 | 5.86 ± 2.21 | 6.00 | 2.00 | 9.00 |
| IBP10 | 4.93 ± 1.69 | 5.00 | 3.00 | 8.00 | 5.29 ± 2.58 | 5.50 | 1.00 | 9.00 |
| IBP10_MML | 3.71 ± 1.33 | 3.50 | 2.00 | 6.00 | 5.93 ± 2.16 | 6.00 | 2.00 | 9.00 |
| IBP40 | 5.36 ± 1.50 | 5.00 | 3.00 | 8.00 | 5.00 ± 2.29 | 5.00 | 1.00 | 9.00 |
| WN | 4.57 ± 1.45 | 5.00 | 2.00 | 7.00 | 5.21 ± 2.33 | 5.00 | 2.00 | 9.00 |
| WN10 | 5.36 ± 1.50 | 5.00 | 3.00 | 8.00 | 4.50 ± 2.21 | 4.00 | 1.00 | 9.00 |
| WN40 | 5.57 ± 2.34 | 6.00 | 1.00 | 9.00 | 3.79 ± 2.22 | 3.00 | 1.00 | 9.00 |
| **Tonal** |  |  |  |  |  |  |  |  |
| IBP | 5.20 ± 1.42 | 5.00 | 3.00 | 7.00 | 4.53 ± 1.81 | 4.00 | 2.00 | 8.00 |
| IBP10 | 5.40 ± 1.64 | 5.00 | 2.00 | 7.00 | 4.73 ± 2.12 | 4.00 | 2.00 | 9.00 |
| IBP10_MML | 4.47 ± 1.96 | 5.00 | .00 | 8.00 | 5.40 ± 1.88 | 5.00 | 3.00 | 9.00 |
| IBP40 | 5.93 ± 1.53 | 6.00 | 2.00 | 8.00 | 4.20 ± 2.11 | 3.00 | 1.00 | 8.00 |
| WN | 5.27 ± 2.22 | 5.00 | 1.00 | 8.00 | 4.87 ± 2.88 | 5.00 | 1.00 | 9.00 |
| WN10 | 5.47 ± 1.60 | 5.00 | 3.00 | 9.00 | 4.27 ± 1.91 | 4.00 | 1.00 | 7.00 |
| WN40 | 5.53 ± 1.41 | 5.00 | 3.00 | 8.00 | 4.13 ± 1.60 | 4.00 | 2.00 | 7.00 |

**Table S4:** Model fitting - arousal & valence. df = degrees of freedom; AIC = Akaike Information Criterion; BIC = Bayesian Information Criterion; logLik = log-likelihood; LRT = Likelihood Ratio Test

| Model | R<sup>2</sup> (marginal) | R<sup>2</sup> (conditional) | df | AIC | BIC | logLIK | LRT | p |
| --- | --- | --- | --- | --- | --- | --- | --- | --- |
| **Arousal** |  |  |  |  |  |  |  |  |
| Intercept only: response ~ 1 + (1\|id) | .00 | .33 | 3 | 776.28 | 786.22 | -385.14 |  |  |
| Fitted model: response ~ condition + (1\|id) | .09 | .42 | 9 | 759.72 | 789.54 | -370.86 | 28.56 | <.01 |
| **Valence** |  |  |  |  |  |  |  |  |
| Intercept only: response ~ 1 + (1\|id) | .00 | .37 | 3 | 858.16 | 868.10 | -426.08 |  |  |
| Fitted model: response ~ condition + (1\|id) | .06 | .42 | 9 | 851.69 | 881.51 | -416.84 | 18.48 | <.01 |

**Figure S1:**
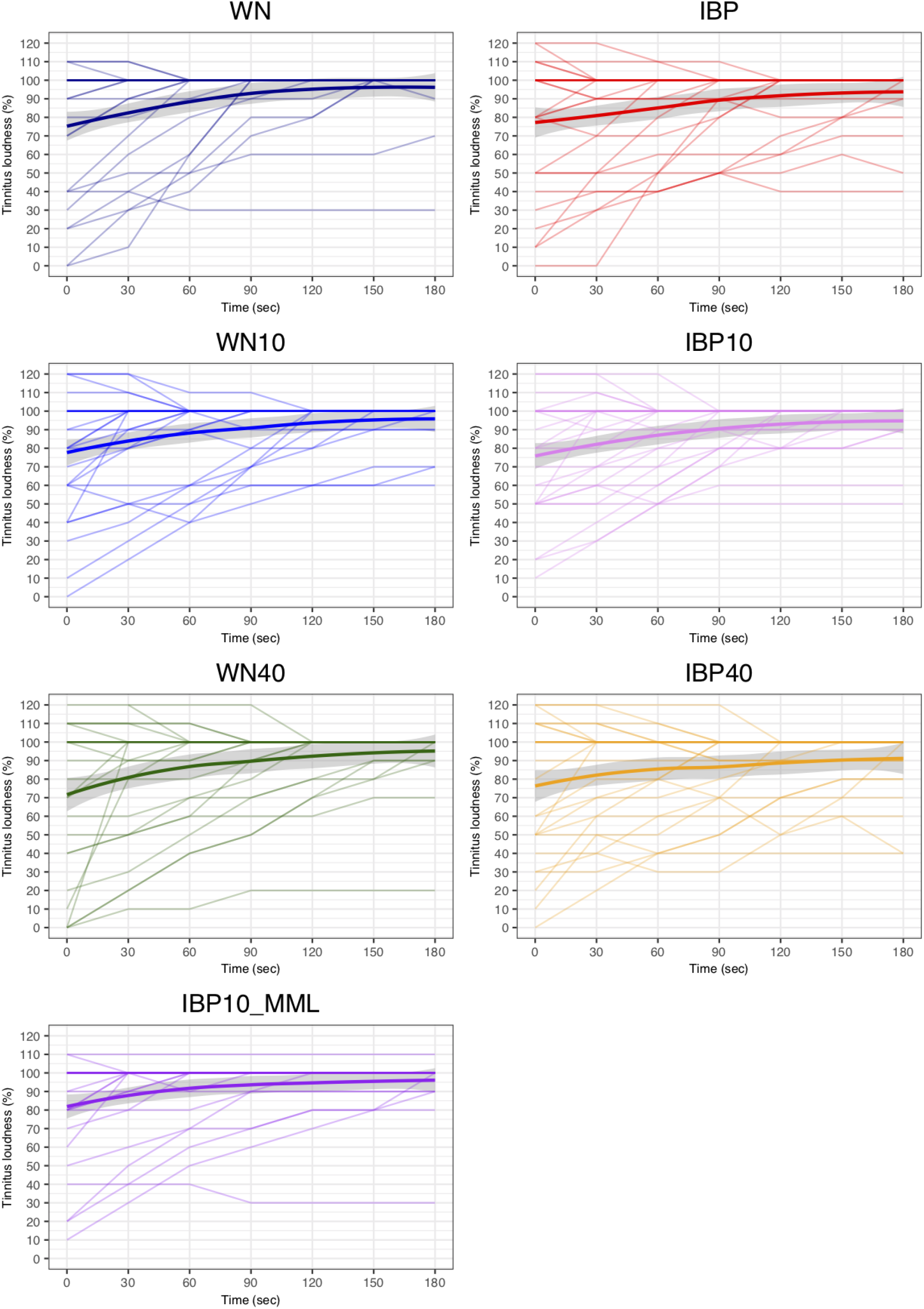
Tinnitus loudness time curve - single patient response. Tinnitus loudness ratings are illustrated on a single participant level for all rating timepoints separated for each stimuli. Thick lines show the mean tinnitus loudness (%) for each stimulus, standard deviations are illustrated as grey ribbons.

## References

Baguley, D., McFerran, D., and Hall, D. (2013). Tinnitus. The Lancet, 382(9904):1600–1607.

Basile, C., Fournier, P., Hutchins, S., and Hébert, S. (2013). Psychoacoustic Assessment to Improve Tinnitus Diagnosis. PLoS ONE, 8(12):e82995.

Bradley, M. M. and Lang, P. J. (1994). Measuring emotion: The self-assessment manikin and the semantic differential. Journal of Behavior Therapy and Experimental Psychiatry, 25(1):49–59.

Eggermont, J. J. (2007). Pathophysiology of tinnitus. Prog. Brain Res., 166:19–35.

Eggermont, J. J. and Roberts, L. E. (2012). The Neuroscience of Tinnitus: Understanding Abnormal and Normal Auditory Perception. Frontiers in Systems Neuroscience, 6.

Eggermont, J. J. and Tass, P. A. (2015). Maladaptive Neural Synchrony in Tinnitus: Origin and Restoration. Frontiers in Neurology, 6.

Erlandsson, S. and Dauman, N. (2013). Categorization of tinnitus in view of history and medical discourse. International Journal of Qualitative Studies on Health and Well-being, 8(1):23530.

Farhadi, M., Mahmoudian, S., Saddadi, F., Karimian, A. R., Mirzaee, M., Ahmadizadeh, M., Ghasemikian, K., Gholami, S., Ghoreyshi, E., Beyty, S., Shamshiri, A., Madani, S., Bakaev, V., Moradkhani, S., and Raeisali, G. (2010). Functional brain abnormalities localized in 55 chronic tinnitus patients: fusion of SPECT coincidence imaging and MRI. J. Cereb. Blood Flow Metab., 30(4):864–870.

Feldmann, H. (1971). Homolateral and contralateral masking of tinnitus by noise-bands and by pure tones. Audiology, 10(3):138–144.

Feldmann, H. (1983). Time Patterns and Related Parameters in Masking of Tinnitus. Acta OtoLaryngologica, 95(5-6):594–598.

Folmer, R. L. (2007). Lateralization of neural activity associated with tinnitus. Neuroradiology, 49(8):689–691; author reply 693–696.

Fournier, P., Cuvillier, A.-F., Gallego, S., Paolino, F., Paolino, M., Quemar, A., Londero, A., and Norena, A. (2018). A New Method for Assessing Masking and Residual Inhibition of Tinnitus. Trends Hear, 22.

Galazyuk, A. V., Longenecker, R. J., Voytenko, S. V., Kristaponyte, I., and Nelson, G. L. (2019). Residual inhibition: From the putative mechanisms to potential tinnitus treatment. Hear. Res., 375:1–13.

Galazyuk, A. V., Voytenko, S. V., and Longenecker, R. J. (2017). Long-Lasting forward Suppression of Spontaneous Firing in Auditory Neurons: Implication to the Residual Inhibition of Tinnitus. Journal of the Association for Research in Otolaryngology, 18(2):343–353.

Goebel, G., Berthold, A., Scheffold, J., and Bläsing, L. (2013). Ein valides Screening-und Evaluationsinstrument zu Erfassung der Hyperakusisbelastung unter Berücksichtigung von Phonophobie und Rekrutiment und Schwerhörigkeit. In Kongreß der Deutschen HNO-Gesellschaft, page 154.

Goebel, G. and Hiller, W. (1994). [The tinnitus questionnaire. A standard instrument for grading the degree of tinnitus. Results of a multicenter study with the tinnitus questionnaire]. HNO, 42(3):166–172.

Hall, D. A., Láinez, M. J., Newman, C. W., Sanchez, T. G., Egler, M., Tennigkeit, F., Koch, M., and Langguth, B. (2011). Treatment options for subjective tinnitus: Self reports from a sample of general practitioners and ENT physicians within Europe and the USA. BMC Health Serv Res, 11:302.

Hallam, R. S., Jakes, S. C., and Hinchcliffe, R. (1988). Cognitive variables in tinnitus annoyance. Br J Clin Psychol, 27 (Pt 3):213–222.

Harrison, X. A., Donaldson, L., Correa-Cano, M. E., Evans, J., Fisher, D. N., Goodwin, C. E., Robinson, B. S., Hodgson, D. J., and Inger, R. (2018). A brief introduction to mixed effects modelling and multi-model inference in ecology. PeerJ, 6:e4794.

Heller, A. J. (2003). Classification and epidemiology of tinnitus. Otolaryngologic Clinics of North America, 36(2):239–248.

Henry, J. A., Roberts, L. E., Ellingson, R. M., and Thielman, E. J. (2013). Computer-Automated Tinnitus Assessment: Noise-Band Matching, Maskability, and Residual Inhibition. Journal of the American Academy of Audiology, 24(6):486–504.

Kleinjung, T., Fischer, B., Langguth, B., Sand, P. G., Hajak, G., Dvorakova, J., and Eichhammer, P. (2007). Validierung einer deutschsprachigen Version des Tinnitus Handicap Inventory. Psychiat Prax, 34(S 1):S140–S142.

Langguth, B., Goodey, R., Azevedo, A., Bjorne, A., Cacace, A., Crocetti, A., Del Bo, L., De Ridder, D., Diges, I., Elbert, T., Flor, H., Herraiz, C., Ganz Sanchez, T., Eichhammer, P., Figueiredo, R., Hajak, G., Kleinjung, T., Landgrebe, M., Londero, A., Lainez, M. J. A., Mazzoli, M., Meikle, M. B., Melcher, J., Rauschecker, J. P., Sand, P. G., Struve, M., Van de Heyning, P., Van Dijk, P., and Vergara, R. (2007). Consensus for tinnitus patient assessment and treatment outcome measurement: Tinnitus Research Initiative meeting, Regensburg, July 2006. Prog. Brain Res., 166:525–536.

Langguth, B., Kreuzer, P. M., Kleinjung, T., and De Ridder, D. (2013). Tinnitus: causes and clinical management. The Lancet Neurology, 12(9):920–930.

Moazami-Goudarzi, M., Michels, L., Weisz, N., and Jeanmonod, D. (2010). Temporo-insular enhancement of EEG low and high frequencies in patients with chronic tinnitus. QEEG study of chronic tinnitus patients. BMC Neuroscience, 11.

Mohan, A., De Ridder, D., and Vanneste, S. (2016). Graph theoretical analysis of brain connectivity in phantom sound perception. Scientific Reports, 6:19683.

Møller, A. R. (2006). Neural plasticity in tinnitus. Prog. Brain Res., 157:365–372.

Nakagawa, S., Johnson, P. C. D., and Schielzeth, H. (2017). The coefficient of determination R2 and intra-class correlation coefficient from generalized linear mixed-effects models revisited and expanded. J R Soc Interface, 14(134).

Neff, P., Michels, J., Meyer, M., Schecklmann, M., Langguth, B., and Schlee, W. (2017). 10 Hz Amplitude Modulated Sounds Induce Short-Term Tinnitus Suppression. Frontiers in Aging Neuroscience, 9.

Neff, P., Zielonka, L., Meyer, M., Langguth, B., Schecklmann, M., and Schlee, W. (2019). Comparison of Amplitude Modulated Sounds and Pure Tones at the Tinnitus Frequency: Residual Tinnitus Suppression and Stimulus Evaluation. Trends in Hearing, 23:233121651983384.

Newman, C. W., Wharton, J. A., Shivapuja, B. G., and Jacobson, G. P. (1994). Relationships among Psychoacoustic Judgments, Speech Understanding Ability and Self-Perceived Handicap in Tinnitus Subjects. International journal of audiology, 33(1):47–60.

Norena, A., Micheyl, C., Chéry-Croze, S., and Collet, L. (2002). Psychoacoustic Characterization of the Tinnitus Spectrum: Implications for the Underlying Mechanisms of Tinnitus. Audiology and Neurotology, 7(6):358–369.

Pantev, C., Okamoto, H., and Teismann, H. (2012). Music-induced cortical plasticity and lateral inhibition in the human auditory cortex as foundations for tonal tinnitus treatment. Frontiers in Systems Neuroscience, 6.

Reavis, K. M., Rothholtz, V. S., Tang, Q., Carroll, J. A., Djalilian, H., and Zeng, F.-G. (2012). Temporary Suppression of Tinnitus by Modulated Sounds. J Assoc Res Otolaryngol, 13(4):561–571.

Roberts, L. E. (2007). Residual inhibition. In. Progress in Brain Research, volume 166, pages 487–495. Elsevier.

Roberts, L. E., Moffat, G., Baumann, M., Ward, L. M., and Bosnyak, D. J. (2008). Residual Inhibition Functions Overlap Tinnitus Spectra and the Region of Auditory Threshold Shift. Journal of the Association for Research in Otolaryngology, 9(4):417–435.

Roberts, L. E., Moffat, G., and Bosnyak, D. J. (2006). Residual inhibition functions in relation to tinnitus spectra and auditory threshold shift. Acta Oto-Laryngologica, 126(sup556):27–33.

Schaette, R., Koenig, O., Hornig, D., Gross, M., and Kempter, R. (2010). Acoustic stimulation treatments against tinnitus could be most effective when tinnitus pitch is within the stimulated frequency range. Hearing Research, 269(1-2):95–101.

Schlee, W., Hall, D. A., Canlon, B., Cima, R. F. F., de Kleine, E., Hauck, F., Huber, A., Gallus, S., Kleinjung, T., Kypraios, T., Langguth, B., Lopez-Escamez, J. A., Lugo, A., Meyer, M., Mielczarek, M., Norena, A., Pfiffner, F., Pryss, R. C., Reichert, M., Requena, T., Schecklmann, M., van Dijk, P., van de Heyning, P., Weisz, N., and Cederroth, C. R. (2018). Innovations in Doctoral Training and Research on Tinnitus: The European School on Interdisciplinary Tinnitus Research (ESIT) Perspective. Front Aging Neurosci, 9.

Schlee, W., Hartmann, T., Langguth, B., and Weisz, N. (2009). Abnormal resting-state cortical coupling in chronic tinnitus. BMC Neuroscience, 10(1):11.

Schlee, W., Schecklmann, M., Lehner, A., Kreuzer, P. M., Vielsmeier, V., Poeppl, T. B., and Langguth, B. (2014). Reduced Variability of Auditory Alpha Activity in Chronic Tinnitus. Neural Plasticity, 2014:1–9.

Terry, A. M. P., Jones, D. M., Davis, B. R., and Slater, R. (1983). Parametric Studies of Tinnitus Masking and Residual Inhibition. British Journal of Audiology, 17(4):245–256.

Tyler, R., Stocking, C., Secor, C., and Slattery, W. H. (2014). Amplitude Modulated S-Tones Can Be Superior to Noise for Tinnitus Reduction. American Journal of Audiology, 23(3):303.

Vernon, J. and Fenwick, J. (1984). Identification of Tinnitus: A Plea for Standardization. The Journal of Laryngology & Otology, 98(S9):45–53.

Vernon, J. A. and Meikle, M. B. (2003). Tinnitus: clinical measurement. Otolaryngologic Clinics of North America, 36(2):293–305.

